# Inducible non-human primate models of retinal degeneration for testing end stage therapies and understanding disease mechanisms

**DOI:** 10.1101/2022.12.03.518955

**Authors:** Divya Ail, Diane Nava, In Pyo Hwang, Elena Brazhnikova, Céline Nouvel-Jaillard, Alexandre Dentel, Corentin Joffrois, Lionel Rousseau, Julie Dégardin, Stephane Bertin, José-Alain Sahel, Olivier Goureau, Serge Picaud, Deniz Dalkara

## Abstract

The anatomical differences between the retinas of humans and most animal models pose a challenge for testing novel therapies. Non-human primate (NHP) retina is anatomically closest to the human retina with the presence of a high acuity region called the fovea. However, there is a lack of relevant NHP models for retinal degeneration that can be used for preclinical studies of vision restoration. To address this unmet need we aimed to generate inducible NHP models of photoreceptor degeneration. We generated three cynomolgus macaque models using distinct strategies. We used two genetically targeted strategies using optogenetics and Crispr-Cas9 to ablate specifically rods to mimic rod-cone dystrophy. Additionally, we created an acute model by physical separation of the photoreceptors and retinal pigment epithelium using a polymer patch. Retinal degeneration was evaluated in all three models by in-life exams such as fundus imaging, optical coherence tomography, adaptive optics and electroretinography. In the genetic models we observed punctuate areas of degeneration in the injected area marked by disorganization of outer segments, loss of rod photoreceptors and thinning of the outer nuclear layer. In the acute model, the degeneration was faster and involved both rods and cones. Among the three distinct NHP models, the Crispr-Cas9 based approach was the most advantageous model in view of recapitulating disease specific features and its ease of implementation. The acute model however resulted in the fastest degeneration making it the most relevant model for testing end-stage vision restoration therapies such as stem cell transplantation.

## Introduction

Retinal degeneration (RD) diseases severely compromise the quality of patients’ lives and constitute an economic burden^1^. Extensive research efforts are focused towards developing effective novel therapies for RD such as gene and cell therapies and retinal implants^2^. However, at present, there is a lack of good animal models to test the effectiveness and translatability of these therapies to humans. Rodents are the most commonly used models but, they lack important anatomical features such a cone-rich macula and fovea, that result in functional differences as well as differences in the manifestation of clinical features and the progression of human diseases^3^. The nonhuman primate (NHP) retina on the other hand is anatomically very close to the human retina, including the presence of a cone-rich macula and fovea, color vision and acuity^4,5^. Also NHPs have the added advantage of similarities with humans regarding genetics and immunology, which would make the manifestation of symptoms and progression of the diseases closer to human patients^6^.

For inherited genetic disorders, an ideal model would be one that carries the mutation and shows similar symptoms as in humans, such as the one generated for Parkinson’s disease^7^. However, generating NHP models by germline transmission is a cumbersome task, and is not permitted in most countries due to national ethical laws^8^. Some NHP facility screens have identified naturally occurring mutations for macular degeneration^9^, retinitis pigmentosa^10^, achromatopsia^11^ and Bardet-Biedl syndrome^12^. But maintenance and breeding of such NHP lines is resource-intensive and not desirable due to time and monetary constraints. Some attempts have been made to circumvent these issues by generating inducible models that recapitulate certain disease features. For example, an NHP model for diabetic ocular neovascularization was generated by AAV-mediated delivery of vascular endothelial growth factor (VEGF), resulting in angiogenesis and changes in the outer nuclear layer (ONL) of the retina^13^. Another study generated a model for glaucoma by intracameral injection of microbeads resulting in an increase in intra-ocular pressure – a hallmark of glaucoma^14^. Different studies have attempted to generate models by induction of focal regions of degeneration using a laser^15–18^. Degeneration has also been caused by injection of chemicals such as cobalt chloride^17^, sodium iodate^19^ and sodium nitroprusside^20^. An NHP model for achromatopsia was generated by Crispr-Cas9 mediated deletion of the CNGB3 gene^21^. An inducible NHP model for rod-cone dystrophy was however still lacking.

To address this unmet need, we used three different strategies to generate NHP models for RD. First, an optogenetic strategy was devised, wherein a photoactivatable protein called KillerRed (KR) was used to induce photoreceptor ablation in a spatially and temporally controlled manner. KR is a dimeric red fluorescent protein genetically engineered from the hydrozoan chromoprotein anm2CP which, is a gene analog of green fluorescent protein (GFP). Upon excitation with green light (540-590nm) KR is activated and produces reactive oxygen species (ROS) that result in oxidative stress and eventual cell death^22^. KR mediated cell ablation is versatile and has been used in different animal models such as *C.elegans*^23^, zebrafish^24,25^ and *Xenopus laevis*^26^. Studies have shown KR-mediated elimination of cancer cells in vitro^27^ as well as tumor reduction by KR-mediated photodynamic therapy in mice^28,29^. A study in the mouse retina involving AAV mediated delivery of KR to Müller glial cells showed successful ablation of these cells, leading to retinal structural and functional anomalies upon illumination with green light^30^.

Our second model was generated by Crispr-Cas9 mediated disruption of the Rhodopsin gene in the rod photoreceptors. It consists of a Cas9 protein coupled with a single-stranded guide RNA (sgRNA) which can be specifically designed to target genes of interest. Insertions and deletions (indels) occur when the sgRNA guides the Cas9 to the gene-of-interest^31^. The Crispr-Cas9 system has emerged as a powerful gene-editing technique and is being applied for generation of large animal models for various diseases^32^. Rhodopsin is a G-protein coupled receptor (GPCR) encoded by the *RHO* gene. It is located in the membrane discs of the outer segments of the rod photoreceptors, and is an important component of the phototransduction cascade that converts light into electrical signals. Mutations in the RHO gene have been identified to be responsible for approximately 10% of patients with Retinitis Pigmentosa (RP) and account for up to 40% of the autosomal dominant RP^33,34^. Rodent models such as the P347S mouse^35^ and P23H rats^36^ with mutations in the Rhodopsin gene are available and are routinely used for vision restoration studies^37^.

Our third model involved the surgical placement of a polymer patch between the retinal pigment epithelium (RPE) and the photoreceptors. The RPE is a monolayer of epithelial cells present at the apical end of the photoreceptor outer segments. The photoreceptors depend on the RPE cells for two vital functions – renewal of the outer segments that are shed daily and re-conversion of the visual pigments (opsins) from their 11-*cis* retinal to the *all-trans* retinal form. Thus, a barrier between the RPE and PRs causes a disruption in these vital functions – resulting in stress, accumulation of toxic byproducts, eventual shortening of segments and cell death^38^ as previously observed in studies of epiretinal implants^39^.

## Results

### Strategy 1: Optogenetic strategy and proof-of-concept in rodents

The KillerRed gene was cloned under the control of a human Rhodopsin promoter for specific expression in the rod photoreceptors (Figure 1A). This construct was packaged into AAV8 vector which is known to efficiently transduce photoreceptors in mice^40^, and delivered by bi-lateral subretinal injections to wild-type (WT-C57BL/6J) mouse retina. The expression of KillerRed was tested by imaging the fundus in the red channel and maximal expression was achieved between 6-8 weeks postinjection (Figure 1E). Retinal flatmounts co-stained for Rhodopsin and KR showed that KR was expressed in the segments of the rod photoreceptors as well as the cell bodies (Figure 1B), which was further confirmed by KR immunolabeling on retinal sections (Figure 1I-I’). In the retinal sections colabeling of Rhodopsin with KR is observed (Figure 1B) whereas the short wavelength cone-opsin does not co-stain with KR (Figure 1C) showing that KR is expressed only in rods and not in cones. Hence, rod-specific expression of KR is achieved with our system.

**Figure 1.**
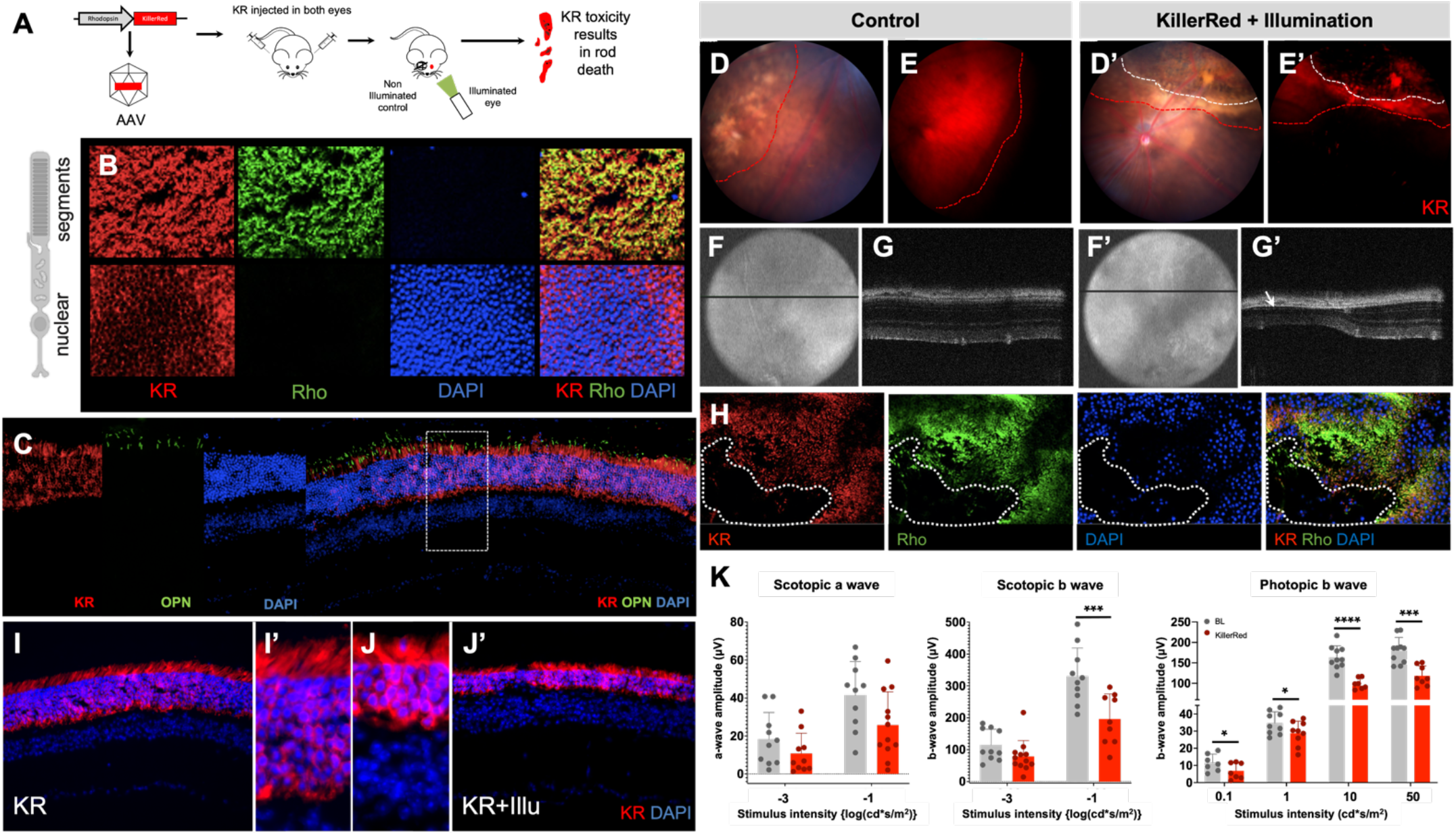
Optogenetic strategy for rod ablation and proof-of-concept in mouse. (A) Schematic representation of the experiment protocol – KillerRed under the control of Rhodopsin promoter is packaged into AAV vectors and delivered by subretinal injection to the retina and activation of KR by light results in toxicity; (B) Immunolabelling of retinal flamounts of KR injected mice with KR (red), Rhodopsin (green), DAPI (blue) and overlay of the 3 channels. Images are acquired at the level of the segments and the ONL; (C) Immunolabelling of retinal section from KR injected eye with KR (red), short-wavelength opsin (green) and DAPI (blue); (D-G’) Fundus images (D,D’), Fundus images in the red channel (E,E’), eye OCT (F,F’) and retinal OCT (G,G’) in KR injected (control) (D-G) and KR injected and illuminated mice (D’-G’). Red dotted line demarcates the injected area and the white dotted line demarcates the area of damage post-illumination; (H) Immunolabelling of retinal flamount from KR injected and illuminated mice. Region of degeneration is demarcated by the dotted white line; (I-J’) KR immunolabelling of retina from KR injected (KR) (I,I’) and KR injected and illuminated mice (KR+Illu) (J,J’); (K) Scotopic a and b wave and Photopic b wave amplitudes shown as a function of stimulus intensity (X axis) from mice before illumination (gray bars) and 4 months post-illumination (red bars), all values are shown as mean ± SD for N = 12 eyes. ONL: Outer Nuclear Layer, OCT: Optical Coherence Tomography.

To induce degeneration, 8 weeks post-injection, one eye was illuminated using an LED light with an average wavelength of 565nm. The other eye was left undilated and covered with a patch to protect from residual light, and served as an injected but non-illuminated control (Figure 1A). The intensity of the light was optimized so as to cause KR activation without light-damage. We used level 4 (corresponding to approximately 5000 lux) in our system (Supp. Figure 1). 4 weeks post-illumination loss of KR expressing rods occurs at the illuminated area. The fundus of these mice shows distinct area of degeneration (yellow dotted line) within the injected area (white dotted line) (Figure 1D’-E’). The optical coherence tomography (OCT) imaging of the both eyes revealed that the different layers of the retina remained intact in the injected control eye (Figure 1F-G), whereas there was severe thinning and complete loss of the outer nuclear layer (ONL) in case of the illuminated eyes (Figure 1F’-G’). Immunolabeling of the flatmount preparations of the KR injected and illuminated retina revealed the loss of rods in the illuminated zone (Figure 1H). KR immunolabelling of retinal sections from the control and illuminated eyes, revealed that there is overall thinning of the ONL even in regions where the rods are not completely eliminated (Figure 1J-J’). Electroretinogram (ERG) was recorded 4 months post-illumination from mice that were injected and illuminated in both eyes (Figure 1K-red bars), and non-injected mice were used as controls (Figure 1K-gray bars). There was a general trend of reduction in the ERG wave amplitudes under both Scotopic and Photopic conditions. This reduction was significant for b wave amplitudes whereas a-wave recordings proved difficult to measure due to high variability (Figure 1K).

### Strategy 1: KR-mediated retinal degeneration in nonhuman primate

We used the same construct of KR under the control of the hRho promoter and packaged it in AAV5, as this serotype has been shown to be most efficient in transducing photoreceptors in NHPs^41^. AAV5-hRho-KR was delivered to both eyes of the NHP (Cynomolgus *macaca fascicularis)* by subretinal injections. Eight weeks post-injection one eye was illuminated, while the other eye was left undilated and covered and served as a control (Figure 2A). The live fundus autofluorescence image showed expression of KR starting from 4 weeks (Figure 2B). KR immunolabeling on retinal flatmounts and sections revealed rod-specific expression with a distinct mosaic pattern observed in primate retinas, confirming we were able to achieve expression only in rods and not in cones of NHP retina (Figure 2C,C’). Post-illumination, the progress of degeneration was monitored by OCT, adaptive optics (AO) and ERG. OCT imaging revealed areas of degeneration in the illuminated eye. The regions of disruption were localized to the ONL as seen in the area demarcated by the yellow dotted lines. (Figure 2D-D”). Heat maps were generated from the injected area post-injection (KR) and 3-months post-illumination (KR+Illu), and the thickness of the entire retina and the ONL was quantified. 3M post-illumination there is thinning of the retina (indicated by black arrows) (Figure 2H). The thickness of the whole retina and the outer retina shows a slight reduction (Figure 2I). Adaptive Optics imaging (AO) gave a clear indication of the degeneration. Small regions-of-interest (ROIs) were selected from the injected and non-injected parts of the retina of both the KR injected but non-illuminated eye (KR-control) and the KR-injected and illuminated eye (KR+Illu) (Figure 2L). Both the density and arrangement of the photoreceptors were affected in the KR+Illu eye (Figure 2K) with an 80% reduction in the density of photoreceptors, and a 27% and 54% increase in dispersion and spacing respectively (Figure 1M). On the other hand, the arrangement and density were not affected in the non-injected areas, although there was a significant effect on the dispersion showing a 38% increase (Figure 2K,2M). The full-field ERG (ffERG) recorded under scotopic and photopic conditions revealed a general trend in reduction of the a and b wave amplitudes in the injected and illuminated eye, but this was not significant (Figure 2F). Rhodopsin immunolabeling revealed the loss of rod photoreceptors and overall thinning of the ONL and retina (Figure 2J). Histological section of the KR injected and illuminated eye revealed that there were broad regions where the ONL was thinner, whereas the adjacent regions showed intact ONL (Figure G1 and G2). On the other hand, there were also regions where the disruptions were more punctuate, localized in smaller clusters showing disorganization of all layers of the retina (Figure 2G3-5). This is also observed by immunolabeling with KR (Figure 2E).

**Figure 2.**
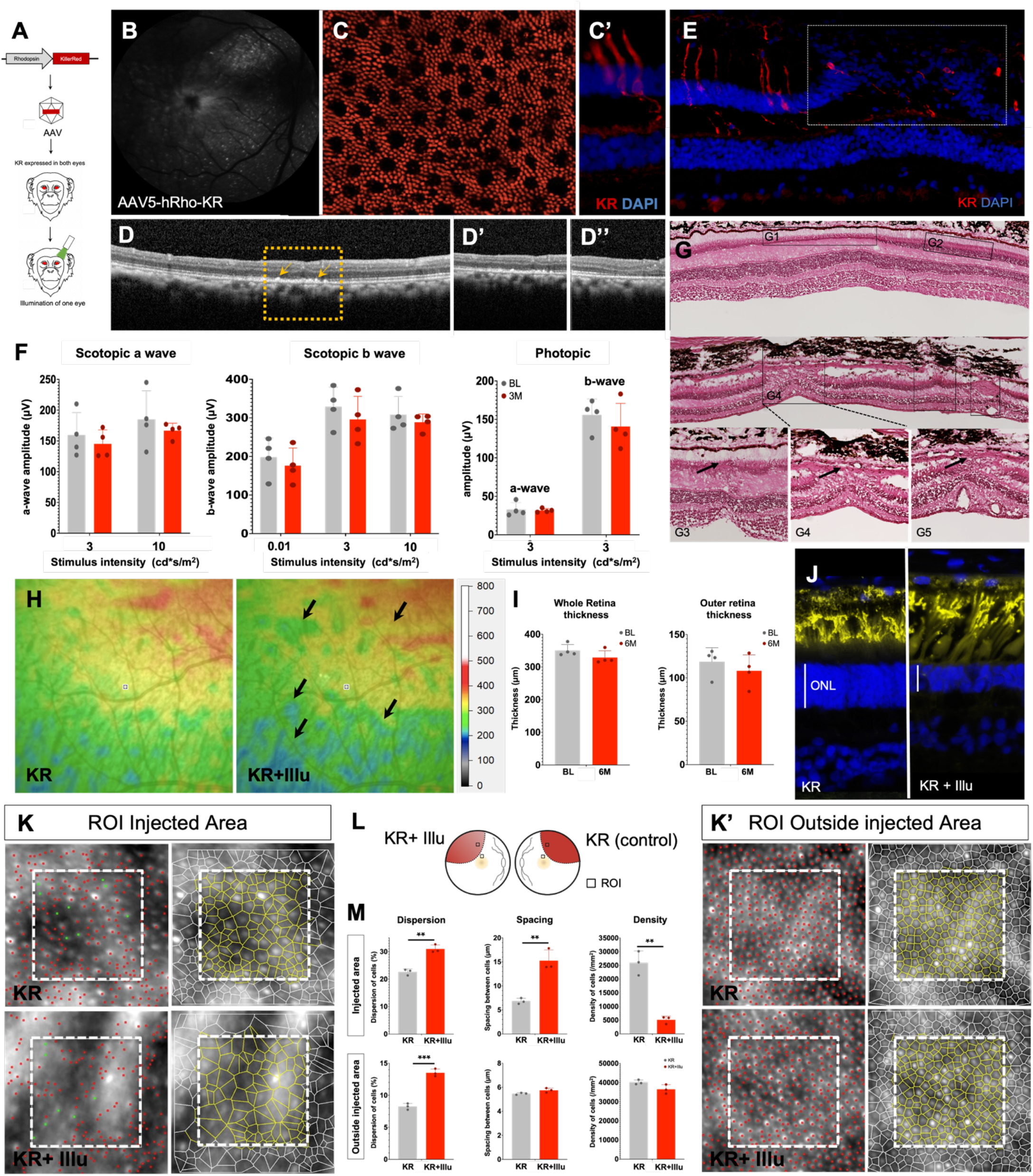
Optogene-mediated retinal degeneration in the nonhuman primate retina. (A) Schematic representation of the experiment protocol – KR under the control of Rhodopsin promoter is packaged into AAV and delivered by subretinal injection to the NHP retina and KR is activated in one eye; (B) Fundus image of KR injected NHP eye; (C,C’) KR immunolabelling in (C) retinal flatmount and (C’) retinal section of KR injected eye showing rod photoreceptor specific expression; (D-D”) OCT images of the KR injected and illuminated eye. Degenerated areas are demarcated by the yellow dotted line and magnified in the corresponding panel (D’); (E) Immunolabelling of KR injected and illuminated eye; (F) Scotopic and Photopic a and b wave amplitudes shown as a function of stimulus intensity (X axis) from NHPs before (BL-gray bars) and post illumination (3M-red bars), all values are shown as mean ± SD for N = 4 NHPs; (G) Histological sections of the KR injected and illuminated eye. Regions of damage are demarcated by the black dotted line and arrows point to degenerated areas of the ONL; (H) Heat maps from the area of injection of eye injected with KR before (KR) and after illumination (KR+Illu). Arrows point towards regions of degeneration and thinner areas of the retina; (I) Thickness of the whole retina and the outer retina 3M post-illumination; (J) Immunolabeling with Rhodopsin (yellow) and DAPI (blue); (K,K’) Adaptive optics images from the region-of-interest (ROI) selected from injected eye (KR) and injected and illuminated eye (KR+Illu) showing the photoreceptors distribution and segmentation in live animals at 6M post-illumination; (L) Schematic representation of the injected and illuminated eye with the ROI used for measurements; (M) quantification of the dispersion, spacing and density of photoreceptors in injected eye (KR-gray bars) and injected and illuminated eye (KR+Illu – red bars). All values are shown as mean ± SD for N = 3 ROIs, * P < 0.05, ** P < 0.001, *** P < 0.0001 with unpaired student’s t-test.

### Strategy 2: Crispr-Cas9 strategy and proof-of-concept in rodents

For the next strategy we used the Crispr-Cas system that has proved to be an efficient tool for gene-editing. The sequence comparison of the exon1 of rhodopsin gene in mouse, macaque and human showed high sequence similarity with some mismatches. Guide RNAs were selected using the Crispor software based on the predicted indel frequency and the off-targets. 4 guide RNAs each specific to mouse, macaque and human sequence were designed and cloned into an expression cassette with the Cas9 under the control of a cytomegalovirus (CMV) promoter. (Supp.Figure 2A). The mouse-specific guides were tested in the photoreceptor cell line – 661W and the sgRNA4 generated the highest indels at 13% (Figure 2B). However, the transfection efficiency of this cell-line was rather low (at 15%) (Supp. Figure 2B’). The macaque and human specific sgRNAs did not result in any indels in 661W cells. Hence, we decided to test the macaque-specific sgRNAs in a monkey COS7 cell line and the human-specific sgRNAs in human HEK cells. The macaque sgRNAs did not result in significant indels (Supp. Figure 2B,B’). The human sgRNA2 at 7% indels was the best performing (Supp. Figure 2C,C’). Interestingly, the human-sgRNA2 sequence was completely identical to the macaque-sgRNA1 and there was a 2bp difference from the mouse-sgRNA4. Hence, we selected mouse-sgRNA4 for the in vivo experiments in mouse and macaque-sgRNA1 for NHP. The construct containing mouse sgRNA4 and CMV-Cas9 was packaged in AAV8 and delivered to the mouse outer retina by subretinal injections (Figure 3B). Two months post-injection the degenerated areas were visible in the fundus images wherein the yellow-dotted line demarcates the severely degenerated areas while the white dotted lines show the injected area (Figure 3C). OCT imaging revealed severe thinning of the outer retina restricted to the injected area 4M post-injection (Figure 3A). This was further confirmed by imaging of histological sections that revealed complete loss of the ONL at 6M post-injection in the injected area (Figure 3D,D’) while the non-injected part of the same eye was intact (Figure 3D,D”). Immunolabelling of retinal sections with Rhodopsin in the control area (Figure 3E,E’) and injected area (Figure 3F,F’) showed complete loss of rods in the injected area. Further, short-wavelength cone-opsin labeling was absent in the injected area compared to the control (Figure 3G,H). Hence, we were able to achieve complete loss of photoreceptors (both rods and cones) 6M post Cas9-Rho injections. The loss of retinal function was tested by ERG by providing increasing flash intensities of light stimulus under scotopic and photopic conditions. There was a general trend in reduction of function in the Cas9-Rho group compared to the Cas9 control. The difference was more significant in the scotopic and photopic b wave amplitudes.

**Figure 3.**
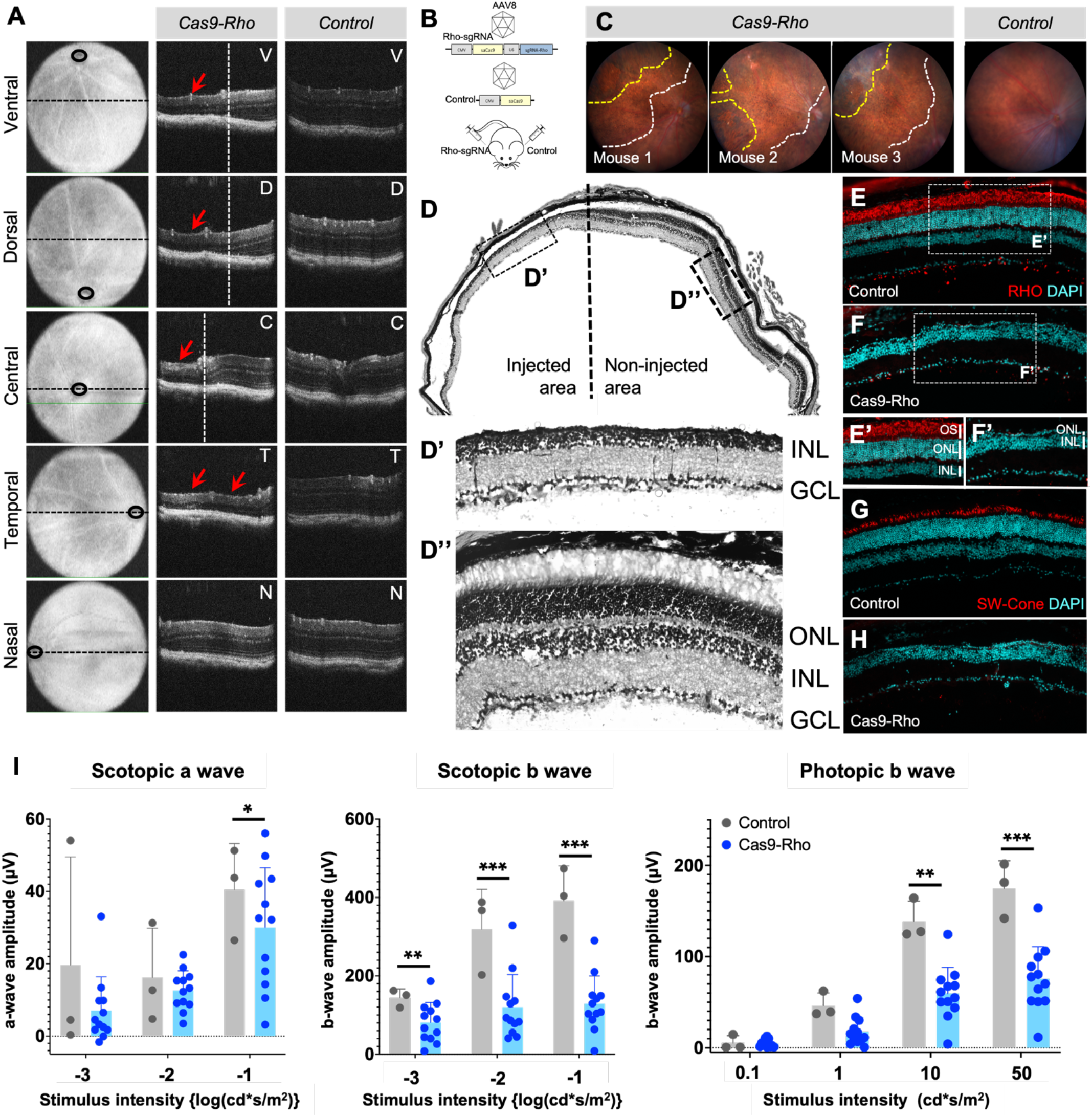
Crispr-Cas9 strategy for rod ablation and proof-of-concept in mouse. (A) OCT images of mouse injected with Cas9 and Rhodopsin sgRNA and control mouse at the ventral (V), dorsal (D), central (C), temporal (T) and nasal (N) regions. Red arrows point to the regions of degeneration; (B) Schematic representation of the experiment protocol in mouse. Cas 9 and guide RNA targeting Rhodopsin under the control of CMV promoter is packaged into AAV vectors and delivered by subretinal injection to the retina; (C) Fundus images of 3 mice post-injection. White dotted line demarcates the injected area and the yellow dotted line demarcates the area of damage; (D) Histology of the eye injected with Cas9-Rho showing the injected and non-injected region. Magnified images of the areas demarcated within the boxes are shown below – (D’) injected area and (D”) non-injected area; (E-H) Immunolabelling of the eye injected with Cas9-Rho (F,H) and the control eye (E,F) with Rhodopsin (E,F) and Short wavelength (SW) cone-opsin (G,H) (in red) and nuclear staining by DAPI (in blue); (I) Scotopic (a and b wave) and Photopic b wave amplitudes shown as a function of stimulus intensity (X-axis) of control (gray bars) and Cas9-Rho injected (blue bars) eyes. Values are shown as mean ± SD for N=3 for control and N=12 for Cas9-Rho eyes and each group was comparted by student’s t test, * P < 0.05, ** P < 0.001, *** P < 0.0001.

### Strategy 2: Crispr-Cas9 mediated retinal degeneration in nonhuman primate

For NHP we developed constructs containing the macaque-specific sgRNA1 targeting the exon1 of rhodopsin gene under the ubiquitous CMV promoter (as used in the Crispr proof-of-concept experiments in mice), or under a rod cell-specific promoter – hRho (as the one used in the KillerRed experiments). For the control eye a Cas9-scrambled guide was used under the control of CMV or hRho promoters. These constructs were packaged into AAV5 vectors and delivered to the adult NHP eyes by subretinal injections (Figure 4A). AO images from the injected area of the control eye (Figure 4B) and the Cas9-Rho eye (Figure 4C) were acquired for analysis and quantification. Both density and arrangement of the photoreceptors were affected in the Cas9-Rho eye (Figure 4C’, C”) compared to the control (Figure 4B’, B”). The density of photoreceptors was 32% lower while the dispersion was 42% higher and spacing was 18% higher compared to the controls. The OCT images 3M post-injection revealed areas of degeneration that were restricted to the outer retinal layers and more specifically to the photoreceptor segments. These ONL disruptions were more severe in the NHP that received Cas9 expressed under a ubiquitous promoter compared to the one that received it with rod cell-specific promoter – hRho. Heat maps generated from the injected areas of the control and Cas9-Rho eyes 3-months post-injection revealed a thinning of the retina (indicated by black arrows) (Figure 4F,G). The thickness of the whole retina and the outer retina was quantified from 3 NHPs and showed a slight reduction 3M post-illumination (Figure 4H). FfERG recordings under scotopic and photopic conditions from the 3 NHPs showed a significant reduction in both the a and b wave amplitudes of the Cas9-Rho group (Figure 4I). Rhodopsin immunolabeling revealed the loss of rod photoreceptors and overall thinning of the ONL and retina (Figure 2J-K’). However, the photoreceptor layer was not completely lost as seen in mice. Histological section of the Cas9-Rho injected eye revealed that there were broad regions where the ONL was thinner, whereas the adjacent regions showed intact ONL (Figure L,L’).

**Figure 4.**
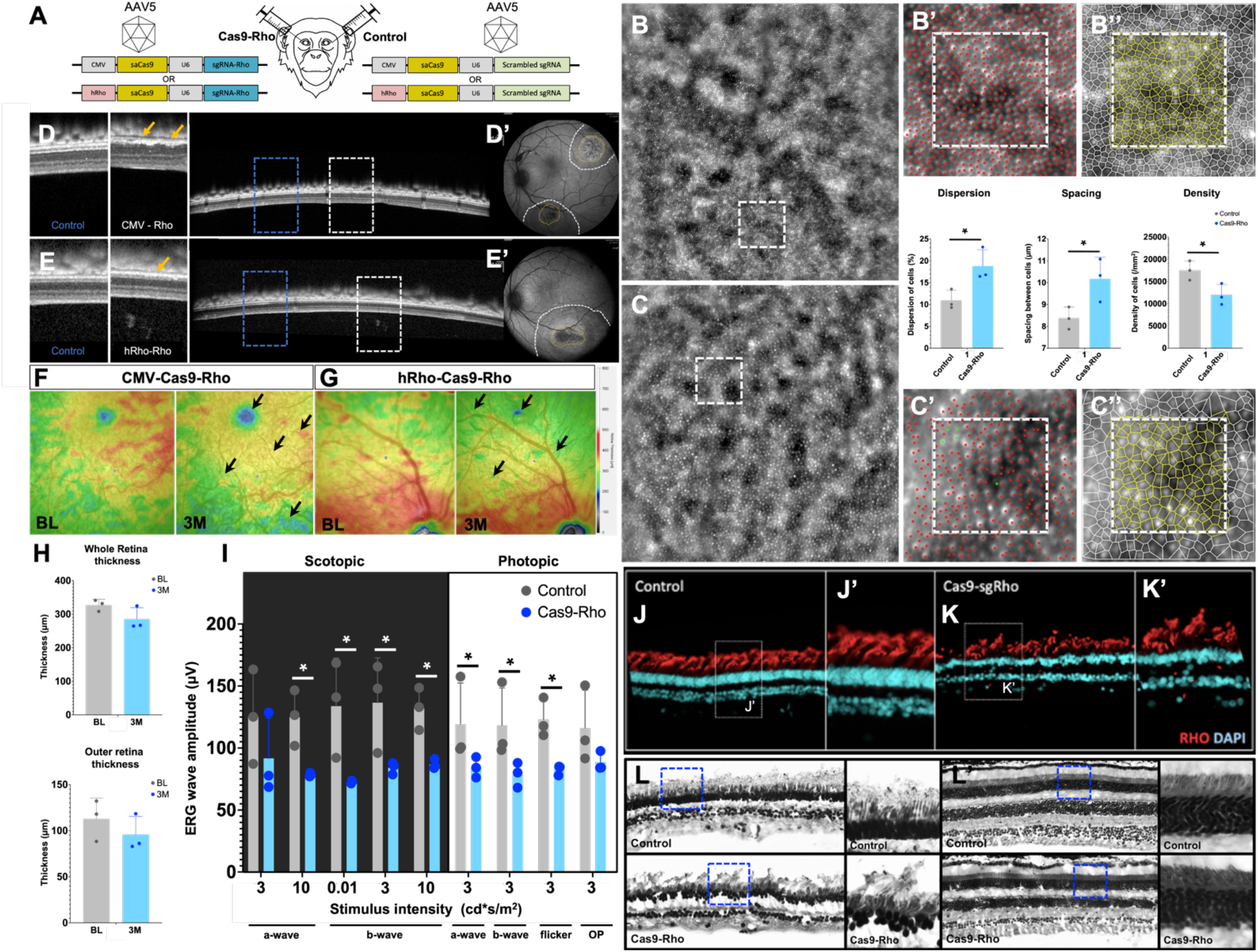
Crispr-Cas9 mediated retinal degeneration in nonhuman primate. (A) Schematic representation of the injection in NHP. Cas9 and guide RNA targeting Rhodopsin under the control of CMV or Rhodopsin promoter is packaged into AAV vectors and delivered by subretinal injection to the retina; (B-C”) Adaptive optics images from the region-of-interest (ROI) (B,C) selected from control eye (B-B”) and Cas9-Rho eye (C-C”) showing the photoreceptors distribution (B’,C’) and segmentation (B”,C”) in live animals at 3M post-injection. Dispersion, Spacing and Density of photoreceptors is quantified in the control eye (gray bars) and Cas9-Rho injected eye (blue bars). All values are shown as mean ± SD for N = 3 ROIs, * P < 0.05 with unpaired student’s t-test. (D-E’) OCT images of NHP eyes injected with Cas9-Rho under the control of an ubiquitous (CMV) (D,D’) or a rod-specific (Rhodopsin) promoter (E,E’). The area demarcated by dotted line and the corresponding magnified images show the unaffected regions (blue boxes as control) and the degenerated areas (white boxes); The injection bleb (white dotted line) and the focal regions of degeneration (yellow dotted line) are demarcated on the eye fundus image (D’,E’); (F,G) Heat maps from CMV-Cas9-Rho injected eye (F) and Rho-Cas9-Rho injected eye at baseline before injection (BL) and 3 months post-injection (3M); (H,H’) Thickness of the whole retina (H) and the outer retina (H’) 3M post-injection; (I) Scotopic (a and b waves), Photopic (a and b waves), flicker ERG and oscillatory potential (OP) amplitudes shown as a function of stimulus intensity (X axis) from control eyes injected with Cas9-scrambled-sgRNA (BL-gray bars) and Cas9-Rho (blue bars), all values are shown as mean ± SD for N = 3 NHPs and each group was comparted by student’s t test, * P < 0.05; (J-K’) Immunolabeling of Rhodopsin in Control (J,J’) and Cas9-Rho (K,K’) eyes; (L,L’) Histological sections of the control and Cas9-Rho eyes from the injected area (L) and a region distal region (L’). Regions of interest are demarcated within blue dotted lines and shown magnified.

### Strategy 3: Physical barrier strategy and proof-of-concept in rodents

In order to have a faster and acute model of degeneration, we envisioned introducing a physical barrier between the RPE and the photoreceptors (PRs). Since the PRs depend on the RPE for their normal functioning and homeostasis, this loss of connection is likely to cause PR stress and eventual death. The proof-of-concept studies were carried out in rats as the mouse retina proved too small for such surgeries (Figure 5A). The patches were made from three different materials – polyimide, polyimide coated with parylene and SU8 and surgically placed between the RPE and PRs. The parylene coating provides a protective layer and allows long-term usage in contact with biological tissues. Both the polyimide (Figure 5C, C’) and polyimide+parylene (Figure 5D,D’) patches were comparable in terms of ease of placement and the level of degeneration achieved. Ideally these patches will be placed for a specific duration of time and when the desired level of photoreceptor loss is achieved that patch will be removed to introduce a therapeutic agent (such as PR transplants) in the degenerated area. In this context, it would be desirable to have a patch that is biodegradable with which we could bypass the patch removal step. The SU8 patches were chosen for their biodegradable properties. However, 2 weeks post-surgery the patches already degraded and caused massive physical damage and inflammation in the surgery location and surrounding areas precluding the use of this material (Figure 5E,E’). Hence polyimide+parylene patches were selected for further experiments in NHPs. The polyimide+parylene patch caused loss of ONL in the region of patch (Figure 5F,G) and overall thinning of the ONL in regions proximal to the patch when compared to control areas distal to the patch (Figure 5H). Immunolabelling with Rhodopsin revealed loss of PRs at the region of the patch compared to the undamaged area distal to the patch of the same eye or the contralateral control eye (Figure 5I). Due to the small size of the eye the patch causes a lot of physical damage during placement (Figure 5G, I). Reducing the dimensions (1.5mm width with 15 μm thickness) further would make them very fragile and difficult to handle. Hence further characterization of these patches in rats was not performed.

**Figure 5.**
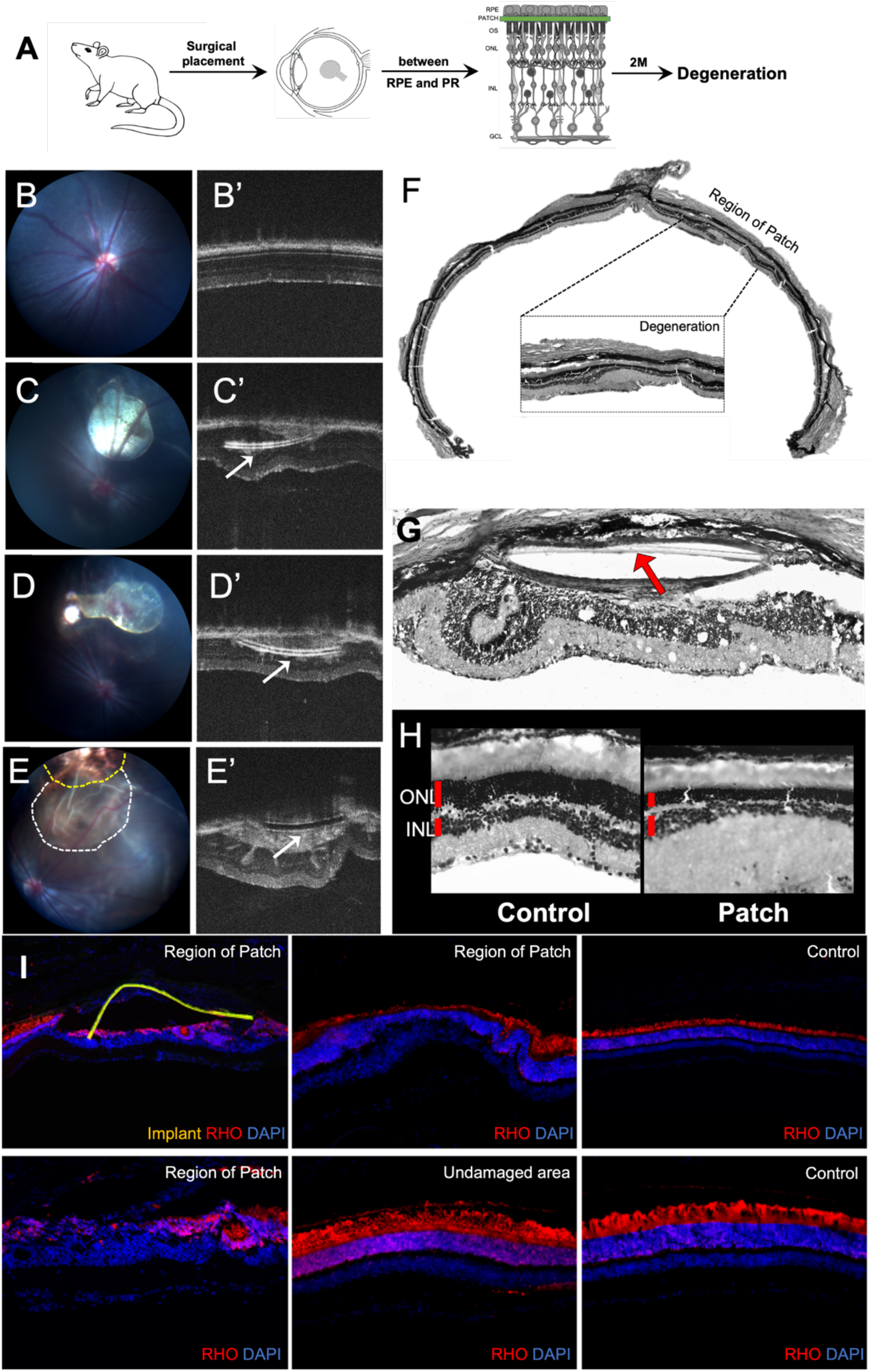
Physical barrier strategy and proof-of-concept in rat. (A) Schematic representation of the experiment protocol – a polymer patch is surgically placed between the RPE and the photoreceptors of rat retinas; (B-E’) Fundus (B-E) and OCT images (B’-E’) acquired before surgery (B,B’) and 4 weeks post-surgery (C’-E’) with polyimide patch (C,C’), parylene coated polyimide patch (C,C’) and SU8 patch (E,E’), arrow points to the patch; (F-G) Histology of the eye with the red arrow pointing towards the polymer patch, (H) magnified images of an area distal from the patch (control) and the area proximal to the patch (Patch); (I-K) Immunolabelling of different regions with Rhodopsin (in red) and nuclear staining by DAPI (in blue). Undamaged area: distal region from the patch, Control: contralateral eye without a patch.

### Strategy 3: Physical barrier mediated retinal degeneration in nonhuman primate

The polyimide+parylene patch was placed surgically between the RPE and PR layer and degeneration was analyzed between one to six months post-surgery (Figure 6A). The patch was placed close to the fovea (blue arrow) (Figure 6B). One-month post-surgery the thickness heat maps showed thinning of the retina close to the patch region as well as the macular region (indicated by arrows) (Figure 6C). The OCT images acquired 1M post-surgery showed that the regions distal to the patch (area1) are unaffected, whereas the ONL was thinner in the region of the patch (area2) and the retinal layers were slightly disorganized in the proximal regions (red arrows) (area3) (Figure 6E). Quantification of thickness revealed that at 1M post-surgery the retina, particularly the outer retina showed some thinning (Figure 6D). AO images from the region with patch and a control region were acquired at and quantified at one- and six- months post-surgery. The density of PRs did not change at one month, but the arrangement was slightly affected as seen by a 1% increase in the dispersion. However, six months post-surgery there was a 26% reduction in the density of PRs, 23% increase in the dispersion and a 14% increase in spacing (Figure 6F). Immunolabeling with Rhodopsin revealed thinner ONL in the region of the patch (Figure 6G2-3) while the adjacent areas remained intact (Figure 6G1,G4). In the region of patch, the ONL is thinner but not completely lost. However, disorganization of the outer retinal layers was apparent (Figure 6H,H’). Comparison of the histological sections showed intact retinal layers and photoreceptors in the control region while there was loss of PRs and thinning of the retina in proximity of the patch (Figure 6I).

**Figure 6.**
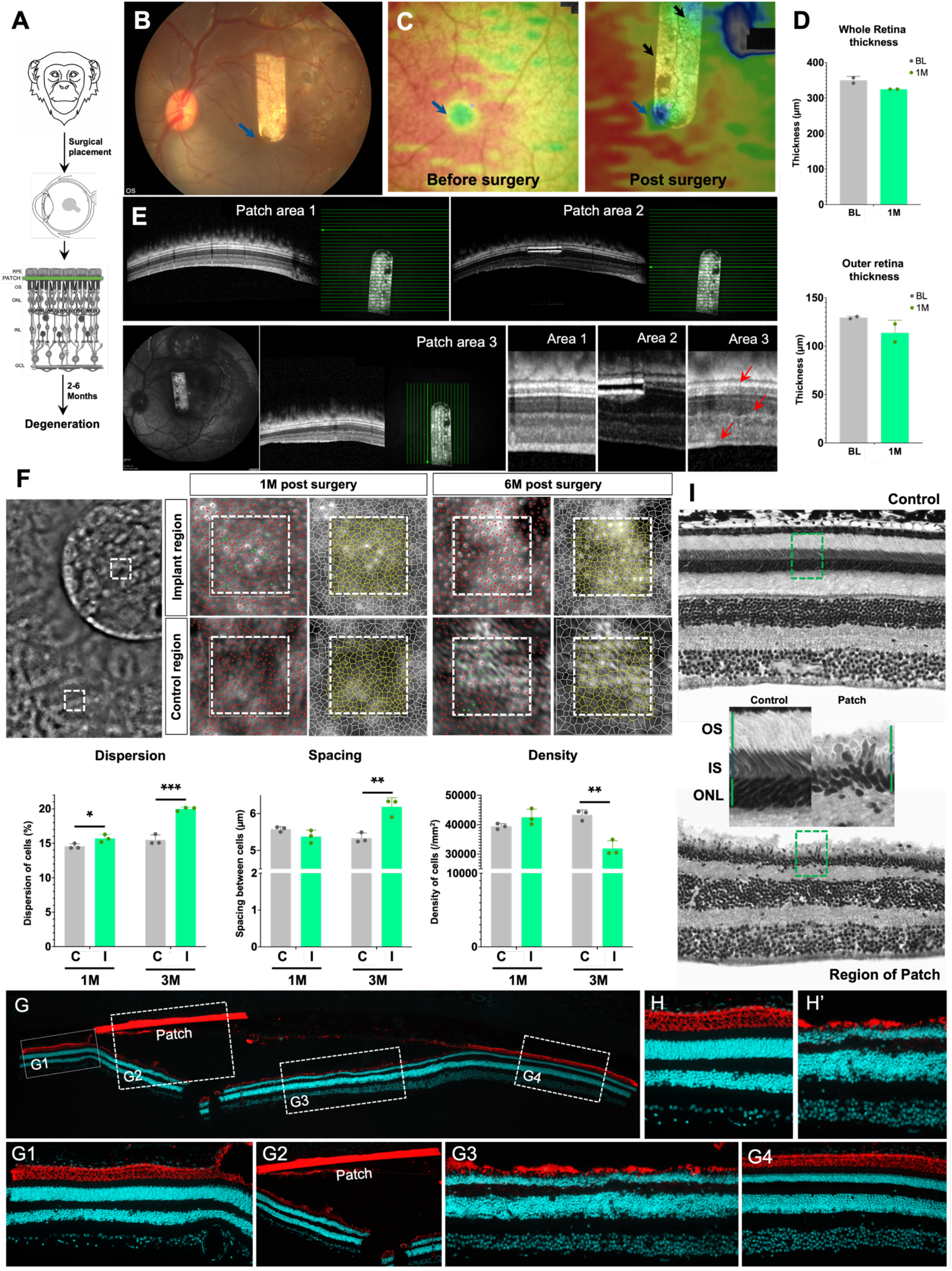
Physical barrier-mediated retinal degeneration in nonhuman primate. (A) Schematic representation of the experiment protocol – a polymer patch is surgically placed between the RPE and the photoreceptors of the NHP retina; (B) Fundus image of the NHP retina showing the placement of the polymer patch in proximity of the fovea; (C) Heat maps from the area of injection of eye injected with the polymer patch before and 1M after surgery, black arrows point towards regions of degeneration and thinner areas of the retina, blue arrow points to the fovea; (D,D’) Thickness of the whole retina (D) and the outer retina (D’) 1M post-injection; (E) OCT images from a region distal from patch (patch area 1), showing cross-section of the patch (patch area 2) and in close proximity of the patch (patch area 3). The bright green lines show the exact region of the OCT images and magnified images from each area are shown in the panels on the right. Red arrows point to punctuate retinal disorganization; (F) Adaptive optics image from the eye with patch showing the region-of-interest (ROI) in the Implant region and control region as white boxes. The photoreceptors distribution and segmentation from the implant and control region is shown 1M and 6M post-surgery. Dispersion, Spacing and Density of photoreceptors is quantified in the control region (gray bars) and implant region (green bars) at 1M and 6M postsurgery. All values are shown as mean ± SD for N = 3 ROIs, * P < 0.05, ** P < 0.001, *** P < 0.0001 with unpaired student’s t-test; (I) Histological sections of the control eye and the eye with patch. Regions of interest are demarcated by the green dotted lines and shown magnified; (G) Immunolabeling with Rhodopsin (in red) and nuclear staining with DAPI (in blue) of the eye with the implant. The different regions-of-interest are magnified in the panels below (G1-4), (H,H’) Immunolabeling of rhodopsin in a distal area (H) and a proximal area (H’) to the implant.

## Methods

### Animals and Ethics statement

Wild-type C57BL6/j mice and Long-Evans Rats were acquired from Janvier Labs and the nonhuman primates (Cynomolgus *Macaca fascicularis)* were acquired from Noveprim, Mauritius. All mice and rats were housed under a 12-h light-dark cycle with food and water ad libitum. The NHPs were screened for pre-existing antibodies against AAV5, and only seronegative animals were used in the study. The NHPs were regularly assessed by clinical observation, monitoring of food consumption and body weight, as well as ophthalmic examination post-injection (fundus imaging, slit lamp examination, OCT and AO). All animal experiments and procedures were ethically approved by the French “Ministère de l’Education, de l’Enseignement Supérieur et de la Recherche” and were carried out according to institutional guidelines in adherence with the National Institutes of Health Guide for the Care and Use of Laboratory Animals as well as the Directive 2010/ 63/EU of the European Parliament.

### Plasmids and AAV production

The KR from the plasmid vector – pKillerRed-mem (Evrogen, Moscow, Russia) was cloned with a human Rhodopsin promoter. The resulting plasmid – hRho-KR was packaged into the AAV8 for injections in mice and into AAV5 for injections in NHPs. Based on the Rhodopsin gene sequence of Mus musculus, Macaca fascicularis and Homo sapiens, we identified all PAM sequences for SaCas9 on the exon1 of the Rhodopsin gene. For each species, the top four sgRNA sequence were selected based on the high on-target score and low off-target score predicted by CRISPOR (http://crispor.tefor.net/). To construct SaCas9 and sgRNA expressing plasmid, the selected sgRNAs were subcloned into pX601-AAV-CMV::NLS-SaCas9-NLS-3xHA-bGHpA;U6::BsaI-sgRNA plasmid (pX601) (Addgene, plasmid 61591 from Feng Zhang). The Cas9 and Cas9-Rho guide containing plasmids were packaged into AAV8 (for injections in mice) and into AAV5 (for NHP injections). Recombinant AAVs were produced using the triple-transfection method on HEK293 cells (ATCC CRL-1573), harvested 24–72 h post transfection and purified by iodixanol gradient ultracentrifugation 42. The 40% iodixanol fraction was collected after a 90 min spin at 354000 g. Concentration and buffer exchange were performed against PBS containing 0.001% Pluronic. AAV vector stocks titers were then determined based on real-time quantitative PCR titration method using ITR primers ^43^ and SYBR Green (Thermo Fischer Scientific).

### Cell transfection and Indel analysis

661W cells (RRID: CVCL_6240), NIH 3T3 (ATCC® CRL-1658™), COS-7 cells (ATCC® CRL-1651™) and HEK 293T cells (ATCC® CRL-3216™) were grown in a 75cm^2^ flask with a culture medium composed of Dulbecco’s Modified Eagle Medium (DMEM), High glucose (Gibco^TM^) supplemented with 10% (vol/vol) of heat-inactivated fetal bovine serum (FBS) (Gibco^TM^) and 1% (vol/vol) of penicillin/streptomycin (Gibco^TM^) in an incubator at 37°C with a 95% O_2_, 5% CO_2_ humidified atmosphere. Cells were passaged using Trypsin-EDTA 0.05% (Gibco^TM^) every three or four days at a split ratio of 1:20. 24h prior the transfection, 8×10^4^ 661W cells were seeded in a 24-well culture plate with DMEM and transfected with the plasmids containing guides using Lipofectamine™ Cells were harvested 72 hours following the transfection and Genomic DNA was then extracted using NucleoSpin® Tissue kit (Macherey-Nagel) following the manufacturer’s recommendation.

For indel analysis, primers were designed to amplify a ~700 bp genomic regions flanking the sgRNA target site with primers annealing at least 200 bp upstream or downstream the cleavage site. PCR products were then purified using NucleoSpin® Gel and PCR Clean-up and sent for Sanger sequencing (Eurofins genomics) using the same forward and reverse primers. Indel values were obtained using the TIDE web tool (https://tide.deskgen.com/) as described previously^44^.

### Subretinal Injections

WT C57BL6/j mice (Janvier Labs) were used in this study and injected at 6 weeks of age. The mice were anesthetized by isofluorane inhalation. Following dilation of the pupils, 1 μl of the AAV was delivered subretinally using a 33-gauge needle. Cynomolgus macaques (Noveprim, Mauritius) were anesthetized with an intramuscular injection of ketamine, 10mg/kg (Imalgene 1000, Merial) and xylazine 0.5mg/kg (Rompun 2%, Bayer). Anesthesia was maintained with an intravenous injection of propofol, 1ml/kg/h (PropoVet Multidose 10mg/ml, Zoetis). Their pupils were dilated and their eyelids were kept open using eyelid speculum. To perform subretinal AAV injections, two 25-gauge vitrectomy ports were set approximately 2 mm posterior to the limbus, one for the endo-illumination probe and the other for the subretinal cannula. A 1-ml Hamilton syringe equipped with a 25-gauge subretinal cannula with a 41-gauge tip was used for the injection. The endoillumination probe and cannula were introduced into the eye. The viral vector solution (100 μl) was injected subretinally to create a bleb on the dorsal side of the fovea. The instruments were then withdrawn. Post-injection eyes received the ointment Fradexam (Labortoire TVM, France) consisting of a corticosteroid (dexamethasone disodium phosphate, 0,76 mg) and an antibiotic (Framycetine sulfate, 3150UI). In case of appearance of any inflammatory signs following the viral injection, the eyes were treated by periocular injection of 12 - 20 mg of Kenacort (Bristol-Myers Squibb) in addition to ointments. In case this treatment was insufficient, systemic (intravenous and/or intramuscular) injections of corticosteroids were provided.

### In vivo imaging (fundus and OCT)

Mice pupils were dilated using 0.5% tropicamide (Mydriaticum, Théa Pharmaceauticals, France) and 5% phenylephrine (Néosynepherine, Europhta, France). The animals were anesthetized by isofluorane (Axience, France) inhalation and the eyes were kept moistened with Lubrithal (Dechra pharmaceuticals) during the course of the experiment. Fundi were monitored and photographed in 2 modalities (bright-field and red fluorescence) using the Micron IV mouse imaging system (Phoenix Research Laboratories, Pleasanton, CA, USA). For Spectral-Domain Optical Coherence Tomography (SD-OCT), the mice were placed in front of the SD-OCT imaging device (Bioptigen 840 nm HHP; Bioptigen, North Carolina, USA). Eyes were kept moisturized with 0.9% NaCl throughout the procedure. Image acquisition was done by using the rectangular scanning protocol consisting with 1000 A-scans (lines) and 100 B-scans (frames) with a 16 frames/B-scan. The images were acquired in the dorsal, ventral, nasal, temporal and central regions of the retina. The images acquired were processed with Fiji software (available at http://fiji.sc/Fiji).

For NHPs, the anesthesia and pupil dilation were carried out as described earlier. Spectralis HRA+OCT system (Heidelberg Engineering, Germany) was used to acquire the fundus images and fluorescent images using the fundus autofluoresence mode (excitation wavelength of 488 nm and barrier filter of 500 nm), as well as the b-scan SD-OCT acquisition.

### Adaptive Optics

The rtx1 (Imagine Eyes, Orsay, France), a compact AO-FIO commercially available device with a lateral resolution of 2μm, was used to image the photoreceptor layer of the NHP. In the rtx1, retinal images are captured by a low-noise high-resolution CCD camera (Manta; Allied Vision, Stadtroda, Germany), while an AO correction system composed of a Shack–Hartmann wavefront sensor (HASO4 first; Imagine Optic, Orsay, France) and a deformable mirror (mirao52-e; Imagine Eyes) works in closed loop to compensate the aberrations introduced by the eye. Retinal images obtained with the rtx1 have a resolution of 2μm over a 4×4-degree field of view. A defocus of +0.50 diopters (corrected by a modified Badal assembly incorporated in the device) was used. NHP were anesthetized during data acquisition because fixation couldn’t be maintained otherwise. Anesthetized animals were laid on the examination table with their head fixed in order to have their eyes aligned with the camera. Because eyes can’t be manually moved during the experiments (otherwise too much astigmatism is induced, avoiding proper imaging), the entire table needed to be positioned to explore intended retinal areas. The investigator (AD) followed the same procedure during each acquisition: (1) locating the fovea with the help of the retinal vascular pattern, and then (2) capturing at least 5 images centered and around the ROI site, as far as technically achievable. The focus (related to the axial depth in μm) was set between ‘0’ and ‘+50’ in order to capture as sharp as possible the photoreceptor layer. NHP were dilated and anesthetized as mentioned above. The acquisition consisted of a series of 40 frames over 2 seconds with an exposure time of 10ms (averaged by the embedded software to produce 1 final image) in a 4 x 4-degree field size, captured at each retinal location mentioned above. Then, the manufacturer commercial software (AODetect 3.0, Imagine Eyes, Orsay, France) was used for cone detection, segmentation, and regularity analyses. In a manually set sampling window of 80 x 80 pixels of each acquired retinal area (ROI), cones were firstly marked semi-automatically (automatic detection by the software, manually adjusted by EB) and the following estimates of density (Voronoi cell density), inter-cell spacing (the mean distance between each cone photoreceptor and its neighbors), and regularity index were obtained. The sampling window for ROI was placed at the closest location that was free from large vessels to avoid vascular artefacts.

### Electroretinogram

Mice were dark-adapted overnight and anesthetized with a mixture of 80 mg/kg ketamine (Imalgene 1000, Merial)and 8 mg/kg xylazine (Rompun 2%, Bayer). Their pupils were dilated as described before, and the cornea was anesthetized by local application of 0.4% oxybuprocaine (Théa Pharmaceauticals, France). Upper and lower lids were retracted to keep the eye open and proptosed. A contact lens-type electrode contacting the cornea through a layer of lubrithal gel (Dechra pharmaceuticals) was used to record the retinal response, with needle electrodes placed on the head and back used as the reference and ground electrodes, respectively. Body temperature was maintained at ~37 °C with a heating pad. The light stimulus was provided by a LED stimulator (ERG Espion from Diagnosys). Responses were amplified and filtered (1 Hz-low and 300 Hz-high cut-off filters) with a 1 channel DC-/AC-amplifier. Dark-adapted (scotopic) responses were measured in darkness, during flash stimulations of 0.001, 0.01 and 0.1 cd.s/m^2^. Photopic ERGs were recorded at stimulations of 0.1,1, 10 and 50 cd.s/m^2^ following a 5-minute light adaptation. Each dark-adapted or photopic ERG response is the mean of five responses from a set of five stimulatory flashes.

For NHPs, the anesthesia and pupil dilation were carried out as described earlier. The RETI-map-animal system (Roland Consult, Germany) was used to record full-field (ffERG) under scotopic and photopic conditions. The scotopic responses were measured after 20 minute of dark adaptation during flash stimulations of 0.0095, 3.0 and 9.49 cd.s/m^2^. Photopic ERGs were recorded at stimulations of 3.0 cd.s/m^2^ following a 10-minute light adaptation. The dark-adapted or photopic ERG responses are a mean of 20-24 responses.

### Illumination

Mice were anesthetized with a mixture of 80 mg/kg ketamine (Imalgene 1000, Merial)and 16 mg/kg xylazine (Rompun 2%, Bayer), and their pupils were dilated as described earlier. NHPs were anesthetized and their pupil dilated as described earlier, and their eyelids were kept open using eyelid speculum. The animals were placed on a heating pad, and a cover slip was placed on each eye after application of lubrithal gel. The eyes were illuminated with green LED light (540-580nm), directed towards the eye at a distance of 10cm, so the eyes received ~5000lux intensity of light. For mice, each eye was exposed to light for a total duration of 3h (in 2 sessions of 1.5h each, 1 week apart) and for NHPs, each eye was exposed for a total duration of 4h (in 2 sessions of 2h each, 1 week apart). Postillumination, the mice eyes were treated with topical application of Ophthalon gel and the NHP eye with an ophthalmic Vitamin A Dulcis ointment (Allergan, Ireland). Whenever, one eye of a mouse was used as non-illuminated control, the eye was not dilated, and was covered with an opaque eye-patch.

### Surgery for placement of patches

All rats were unilaterally implanted at 8 weeks of age. The surgery consisted of placing the patch in the subretinal space in the central region next to the optic nerve as previously described^45^. Briefly, analgesia was provided with subcutaneous injection of buprénorphine (0.05 mg/kg) (Buprecare, Axience) and anesthesia was provided by 5% gaseous induction of isoflurane maintained at 2-3 %. An additional cornea anesthesia was administered using oxybuprocaïne chlorohydrate eye drops. The eye was dilated by application of tropicamide (Mydriaticum 0.5%) solution. The animal was placed on a heating platform at 37°C to maintain body temperature throughout the procedure. A small sclerotomy was performed on the dorsal sclera tangential to the cornea. Sodium chondroitin sulfate – sodium hyaluronate (Viscoat Alcon) gel was injected in the sclerotomy to generate a retinal detachment. The implant was then inserted below the detached retina in the subretinal space in a location adjacent to the optic disk. At the end of surgery, the animal is placed in a recovery chamber at 30°C and postoperative monitoring was conducted with postoperative analgesia.

Nonhuman Primates were anesthetized by an intramuscular injection of ketamine (10 mg/kg) and xylazine (5 mg/kg) and maintained with an intravenous infusion of Propofol 1 ml/kg/h, followed by a local ocular anesthesia (oxybuprocaine chlorhydrate, Thea). Pupillary dilatation was achieved with 0.5% tropicamide eye drops (Mydriaticum) at least 20 - 25 min before intervention. The non-operated eye was protected with an ophthalmic gel (Lubrithal, Dechra) to prevent dehydration. Subretinal implantation was performed using a 3-port pars plana vitrectomy system with cold vitreous cavity irrigation (BBS plus, Alcona) under a stereomicroscope (Lumera 700, Zeiss). The vitrectomy (23 Ga ports) was realized using Alcon Constellation. A bleb for the retinal detachment was then created by subretinal injection of BSS with a subretinal cannula near the macular area. Prior to retinotomy, the appropriate area of the retina was coagulated. A retinotomy of 2 mm was performed using microscissors. The scleral incision of 2 mm was created to introduce the delivery system via the pars plana into the vitreous cavity. The patch was then delivered through the retinotomy and the sclerotomy was finally closed. The retina was attached under PFCL (Perfluorocarbon liquid DECA, DORC). A laser photocoagulation (Vitra Laser, Quantel Medical) was then applied to the area of the retinotomy. After a fluid/air followed by an air/ 20% SF6 gas (Physiol) exchange, the trocars were removed and the additional portion of the 20% SF6 gas was added if needed. The conjunctive was closed with a 8-0 suture and the animal was allowed to awake progressively by intravenous propofol withdrawal.

### Tissue preparation, immunohistochemistry and microscopy

To prepare retinal sections from mice and rats, the eyes were enucleated and fixed in 4% paraformaldehyde (PFA) prepared in phosphate-buffered saline (PBS), overnight. Cornea and lens were removed, and the eyecups were transferred into a tube containing 30% sucrose in phosphate buffered saline (PBS). For NHPs, the eye was fixed in 4% formaldehyde (Sigma-Aldrich) for 4h at RT, followed by dissection and removal of cornea and lens. The eye cup was then transferred into PBS. A 50mm section was cut out from the region of interest, and fixed overnight at 4 ° C. The dissected part was then treated with a sucrose gradient - 10% for 1h, 20% for 1h and 30% overnight at 4 ° C. Eyecups (for mice and rats) and dissected region (for NHP) were then embedded in tissue optimal cutting temperature compound (Microm Microtech, France) and snap-frozen in liquid nitrogen. 12 μ M thick sections were cut on a cryostat (Leica Biosystems), air-dried, and stored at −80° C till further use. To prepare retinal flatmounts, mice eyes were enucleated and incubated for 5 minutes in 2% PFA. After having removed the cornea and lens, the eyecup was cut in a cloverleaf shape by making four incisions. The retina and RPE were dissected from the sclera and placed separately in PBS in a 48-well plate. Dissected retinas and RPEs were post-fixed at room temperature for 1h with 4% PFA. Similar procedure was followed to prepare flatmounts of the dissected region of the NHP retina.

The retinal sections were incubated with blocking solution (3% goat serum + 0.3% Triton-X in PBS) for 2h at room temperature (RT) (10% goat serum was used for blocking of the retinal and RPE flatmounts), washed with PBS, followed by overnight incubation with the primary antibody (listed in Table 2) at 4° C. The following day, the tissue was washed with PBS and incubated with secondary antibody conjugated with Alexa Fluor dyes 488 (emission in the green channel) or 594 (red channel) for 1h at room temperature. DAPI (4’ −6’ -diamino-2-phenylindole, dilactate; Invitrogen) was used for nuclear staining. After PBS washes, the retinal flatmounts or cryosections were mounted onto slides using Vectashield mounting medium (Vector Laboratories, USA) and protected with cover slips. Immunofluorescence was visualized using an Olympus Upright confocal microscope and then analyzed with Fiji software.

## Discussion

The field of ocular gene and cell therapies is experiencing a massive increase in the development and translation of experimental therapies into clinical application^46,47^. However, there is an unmet need for relevant pre-clinical models that can be used for testing these therapies and understanding disease mechanisms. The several available rodent models serve well for proof-of-concept studies but lack comparable size and anatomical features (macula and fovea) that affect the manifestation and progression of the disease features^3^. Here, we successfully generated three distinct animal models that can be used for testing different gene and cell-based therapies and understanding the progression of disease.

Our main aim was to generate a model for rod-cone dystrophy which are primarily caused by mutations in rod photoreceptor genes^48^. One of the important features of rod-cone dystrophies is that the rods are lost first followed by cone loss since the cones depend on rods for survival factors^49–51^. The adaptive optics (AO) imaging modality was particularly important to test this as it quantifies the cone photoreceptors. And in each of our models about three to six months post-induction of degeneration we observe a reduction in the density of the cones, with disruptions in their spatial distribution. The ‘KillerRed model’ mimics this by causing rod cell death by oxidative stress and a reduction in cones can be observed eight to ten months post-induction similar to the sequential events that occur in patients with rod-cone dystrophy. The ‘ Crispr-Cas9 model’ is genetically representative of a common human rod-cone dystrophy as with this technique we specifically ablate a frequently mutated gene in rods by editing the rhodopsin gene similar to the RHO mutation in human disease. With this strategy reduction in the number of cones is observed three months post-injection. In the ‘Patch model’ we do not specifically target the rods, but a small area of the retina with the intention of clearing both rods and cones in the region. This resulted in a reduction in cones after six months. Here the cones are not lost as a consequence of rod death, but cones and rods are lost simultaneously. In the KR model there is an increase in the dispersion of cones outside the injected area in the macular region which is similar to the dispersion observed in the Patch model at one-month post-surgery. This suggests that there is some lateral effect outside the injected area and probably some cell death and cone loss will be observed in the macular and foveal region with time.

The KR model allows spatial control as the degeneration will be restricted to the injected area, as well as only rods will be lost specifically due to the use of a rod-specific promoter. It further permits temporal control as the damage will be initiated only after exposure to light. In both mice and NHP we observe KR expression for up to twelve months post-injection (data not shown). Due to this unique feature many macaques can be injected with KR and when required for testing, the eye can be exposed to light. For light exposure we used an LED light at approximately 565nm at a specified distance from the eye, specific intensity and exposure-time. This can be further optimized to achieve better temporal control by modulating the duration of exposure and/or changing the distance, intensity and type of LED used to cause focal or dispersed degeneration (depending on the requirement).

In the Crispr model we made an important observation for in-vitro testing of guides. While designing and testing the guides for the Crispr model we used the retinal cell line - 661W that was derived from retinal tumors of transgenic mice^52^. We, did not achieve a high transfection efficiency of this line (approximately 15%) but the guides designed for mouse version of rhodopsin resulted in indels (highest was nearly 12%). As expected, when we tested the macaque guides in a monkey cell-line (COS7) and the human guides in a human cell-line (HEK) we were able to generate indels prior to in vivo testing. Degeneration of the retina occurs in stages and the chosen vision restoration method largely depends on the type and stage of the disease. In the earlier stages, when the retinal cells are still present neuroprotective factors can be provided to slow down the progression of the disease and gene replacement or editing can be used to compensate the effect of gene mutations. On the other hand, in later stages when the retinal cells are mostly lost, cell replacement therapy and placement of electrical implants could help in restoring some useful vision. Additionally, specific therapies can control symptoms that arise from perturbed cellular mechanisms such as oxidative stress, inflammation or neovascularization. Our models can be used for testing a wide range of these therapeutic strategies. There are many neuroprotective factors that have been tested in rodents and shown to slow down or prevent degeneration such as bile acids, dopamine, steroid hormones and neurotrophic factors (GDNF: glial-derived neutotrophic factor, BDNF: Brain-derived neurotrophic factor, CNTF: ciliary nerve trophic factor, NGF: nerve growth factor etc.)^54^. All three models can be used for testing such factors. One potent factor known as RdCVF (rod-derived cone viability factor) is secreted by rods and has been shown to be responsible for the eventual cone death after rod loss^50,55^. The neuroprotective effect of Rdcvf can be tested in the KR and Crispr models, since, these models specifically cause rod death and by adaptive optics we were able to show that the cone death follows. The three models can also be used for testing specific gene therapies provided the cell type where the gene is to be expressed is still present. However, repeat injections to the same area can be technically challenging and undesirable due to the possibility of surgical damage and potential inflammation caused by anti-AAV immune responses^56,57^. To circumvent this novel AAV variants, non-viral vectors or intravitreal modes of delivery may be considered.

All three models can be used for testing mutation independent therapies for late-stage degeneration such as cell transplantation and artificial retinal stimulation by electrical prosthesis or optogenetic therapies. Generation of photoreceptors from induced pluripotent stem cells is well-established now^58,59^. However, attempts to transplant these cells into rodents require strong immunosuppression to avoid graft rejection of human cells in rodents. This needs to be taken into account while designing experiments alongside other factors such as material transfer and lack of proper integration^60,61^. But how the transplanted cells will behave in NHP is not known and the models developed here can be useful to examine the effects of this intervention. The recent clinical success with a patient who had his vision restored following optogenetic therapy^62^ is likely to increase the need for preclinical testing of such therapies. Such therapies can be tested in both the Crispr model and the Patch model by expressing the optogene either in the remaining dormant cones or other cell types of the retina (such as ganglion cells and bipolar cells). The KR model can also be considered for such therapies as well as for tests of retinal prosthesis. However, the retinas response to two surgeries (placing the patch and then replacing it by a retinal implant) is still to be tested.

Although rod-cone dystrophies are inherited conditions, non-genetic factors such as oxidative stress and inflammation are important common mechanisms involved in their pathogenesis and progression^63^. The retina is a metabolically demanding tissue with a high oxygen consumption rate. Additionally, the opsin molecules inside the photoreceptors are being constantly subjected to light and oxidative stress, which further results in accumulation of ROS (reactive oxygen species) in RPE. Oxidative stress can result in activation of specific genes, transcription factors or microRNAs that alter key pathways to further alter cellular response such as microglia activation, gliosis and cell death^64^. In our KR model, light-induced activation of KR causes degeneration by oxidative stress and hence this model provides an opportunity to study molecular mechanisms involved in oxidative stress associated with retinal degeneration. In our models we have also observed gliosis marked by microglia activation and migration, and astrocyte and muller cell activation (data not shown). Hence, immune responses and aspects of inflammatory responses can also be studied in these models.

In conclusion we have addressed an important unmet need in the field and propose three preclinical models that cause degeneration by distinct mechanisms following different timelines corresponding to the various needs of translational researchers working in the field of retinal disease. We have used multiple imaging modalities to monitor the progression of the degeneration in live animals which can be further used to monitor the improvement after therapy. Obtaining significant data with NHP studies implies high cost, time and efforts and by investing these in our models we have secured robust and relevant data that will make these models suitable for testing various mutation dependent and independent therapies.

## Acknowledgements

This work was supported by ERC Starting Grant (REGENETHER 639888), the National Institutes of Health NIH / NEI Grant N°1R21EY031117-01, the Institut National de la Santé et de la Recherche Médicale (INSERM), Sorbonne Université, The Foundation Fighting Blind- ness, Agence National de Recherche (ANR) RHU Light4Deaf, *LabEx LIFESENSES (ANR-10-LABX-65), and *IHU FOReSIGHT (ANR-18-IAHU-01). The authors would like to thank Melissa Desrosiers and Camille Robert for their technical assistance with the production of plasmids and viral vectors. We would like to thank Patrick Flament and MirCEN for assistance with all NHP experiments. The authors also extend their thanks to Cardillia-Joe Simon, Duohao Ren and Claire Petr for assistance with experimental work. The schematic illustrations were created on BioRender.com.

---

**Table 1.**
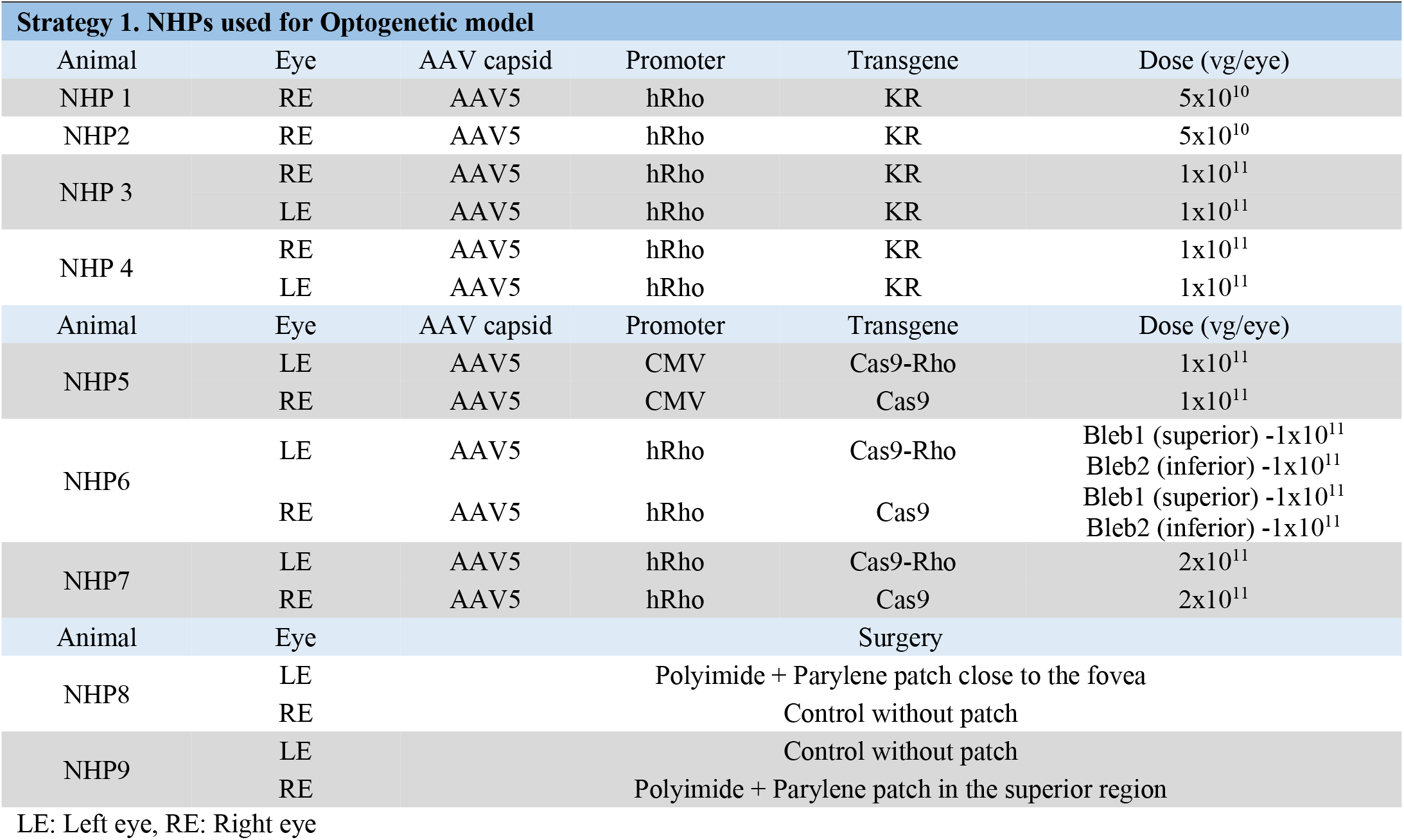
Summary of injections and surgeries in nonhuman primates

**Supplementary figure 1.**
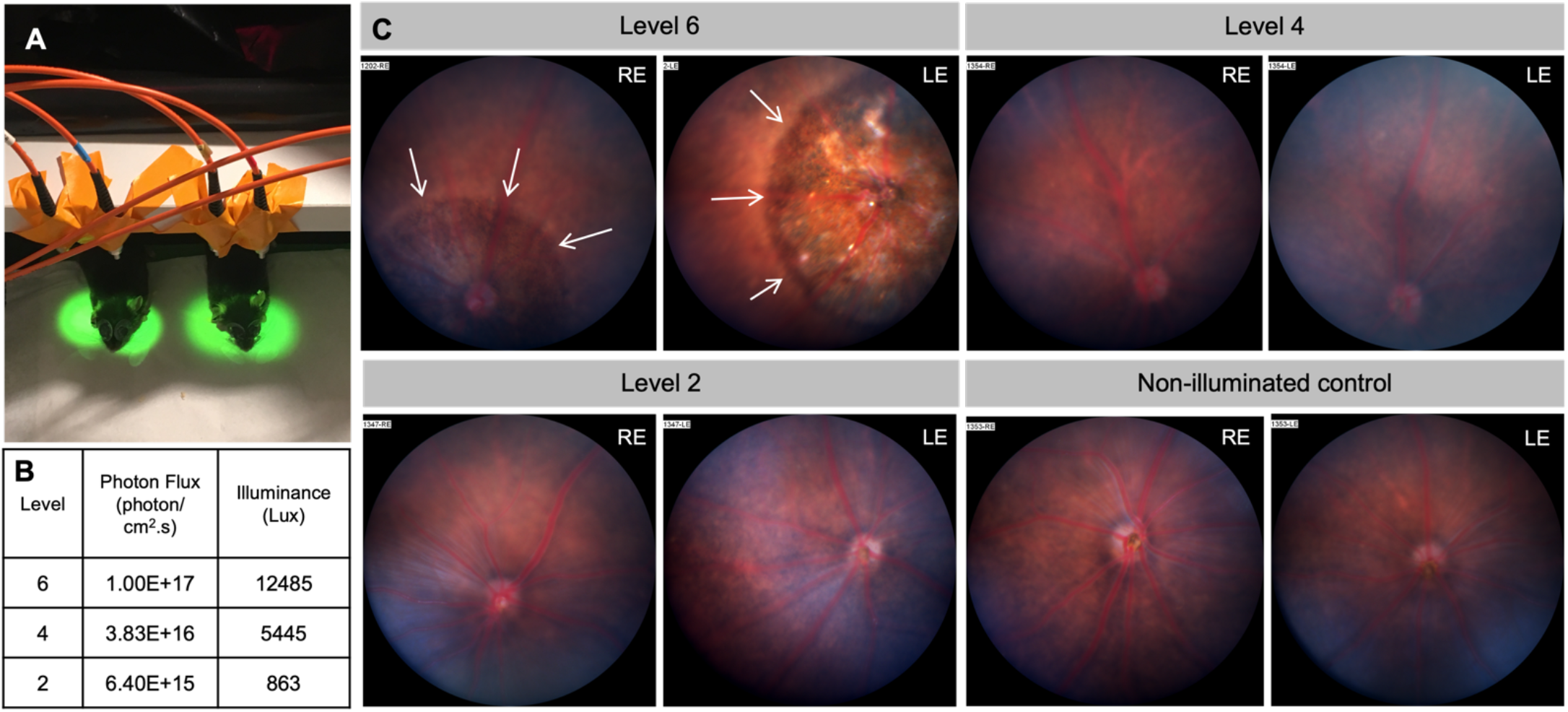
Optimization of the light intensity for KillerRed activation. (A) Set-up of the 565nm LED lights used for activation of KillerRed; (B) Table showing the intensity levels in the apparatus and the corresponding values in Photon flux and luminance; (C) Exposure of mice eyes to Intensity levels 6, 4, 2 and non-illuminated controls. Arrows point to regions of damage. LE: Left eye, RE: Right eye.

**Supplementary figure 2.**
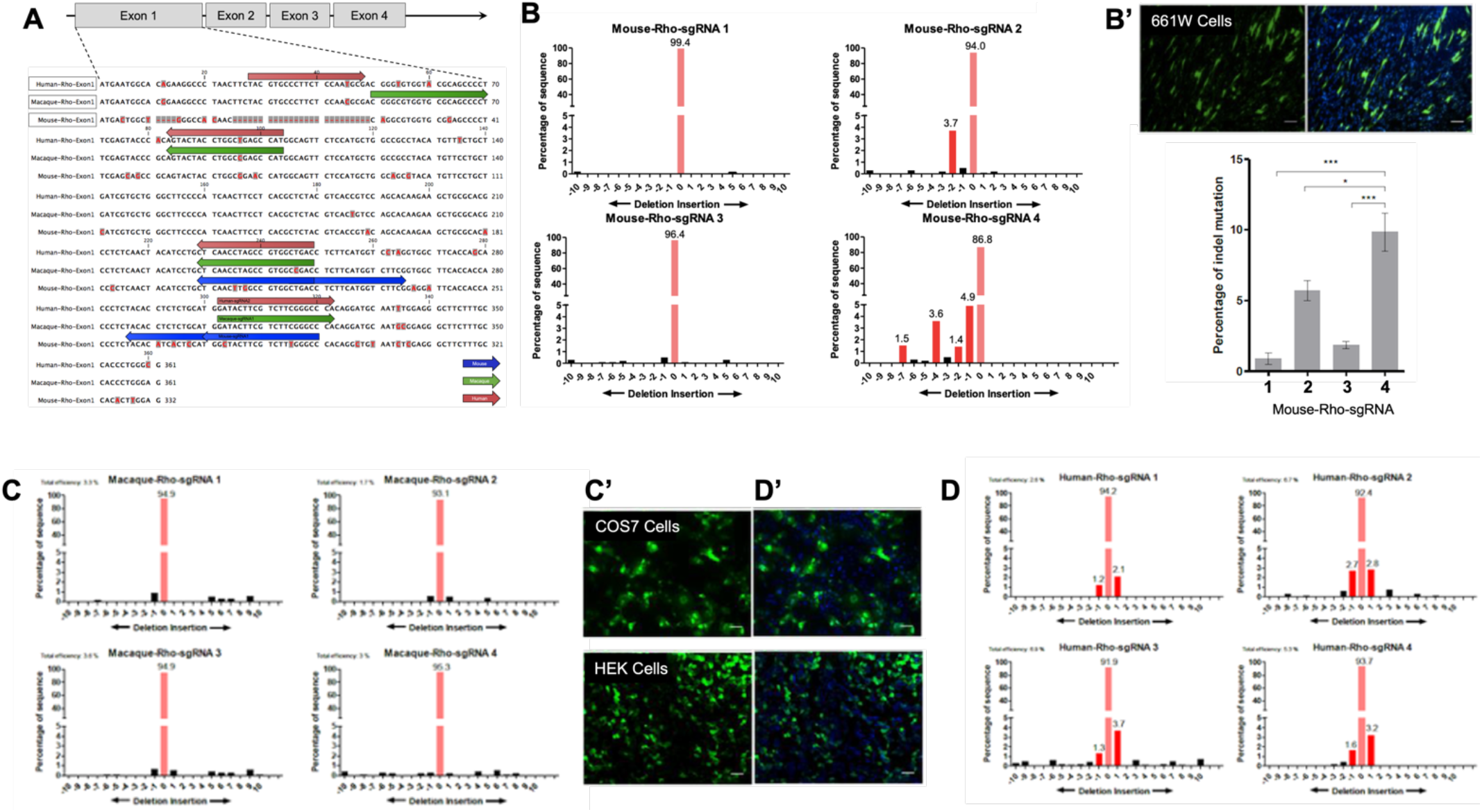
Design and selection of Rhodopsin specific guide RNAs. (A) Schematic of the first 4 exons of the rhodopsin gene showing the sequence of exon1 in mouse, macaque and human. The bases that do not match are highlighted. The sites targeted by guide RNAs designed for mouse (in blue), macaque (in green) and human (in pink) are shown as arrows above the sequence; (B) Indel analysis of the 4 mouse-specific guides tested in the photoreceptor cell line-661W; (B’) Transfection efficiency tested by GFP in 661W cells, (B”) Comparison of the percentage of Indels caused by the 4 guides; (C) Indel analysis of the 4 macaque-specific guides tested in the COS7 cells; (C’) Transfection efficiency tested by GFP in COS7 cells; (D) Indel analysis of the 4 human-specific guides tested in the HEK cells; (B’) Transfection efficiency tested by GFP in HEK cells.

